# Scientific societies advancing STEM workforce diversity: Lessons and Outcomes from the Minorities Affairs Committee of the American Society for Cell Biology

**DOI:** 10.1101/794818

**Authors:** Verónica A. Segarra, Sydella Blatch, Michael Boyce, Franklin Carrero-Martinez, Renato J. Aguilera, Michael J. Leibowitz, MariaElena Zavala, Latanya Hammonds-Odie, Ashanti Edwards

## Abstract

Promoting diversity and inclusiveness in the STEM academic workforce remains a key challenge and national priority. Scientific societies can play a significant role in this process through the creation and implementation of programs to foster STEM academic workforce diversification, and by providing mentoring and skills development training that empower scientists from under-represented minority (URM) backgrounds to succeed in their communities of practice. In this article, we provide examples of challenges met by scientific societies in these areas and present data from the American Society for Cell Biology, highlighting the benefits received by trainees through long-term engagement with its programs. The success of these initiatives illustrates the impact of discipline-specific programming by scientific societies in supporting the development of URM scientists and an increasingly diverse and inclusive academic STEM community.

## INTRODUCTION

The effort to create a diverse and inclusive academic biomedical workforce in the United States continues to face significant challenges. At present, under-represented minority (URM) scientists constitute only 3-4% of medical school basic science tenure-track faculty in Association of American Medical Colleges member institutions, a disproportionately small group in comparison to the ~32% of the U.S. population that are URM (1). Another disparity exists with regard to gender, as women have earned the majority of biomedical Ph.D. degrees from U.S. institutions since 2008, yet represent only one-third of NIH-funded principal investigators (2, 3). While URM and well-represented (WR) trainees are equally likely to matriculate in and complete doctoral programs and to obtain postdoctoral positions, recent analyses indicate that two major bottlenecks exist on the path to becoming faculty: URMs exhibit higher attrition rates during both undergraduate education and in the transition from postdoctoral to junior faculty positions (1, 4). Importantly, although the number of URM Ph.D. graduates increased 9.3-fold between 1980 and 2013, this did not translate into greater representation at the faculty level (4). These observations suggest that merely increasing the total number of URM Ph.D. recipients is unlikely to have a major impact on the proportion of URM biomedical faculty, a conclusion supported by computer modeling of the academic biomedical training and hiring landscape (4). Interventions will likely be necessary across the career trajectory – and particularly at the postdoc-to-faculty transition – to improve representation of URM scientists in the professoriate (1, 4). More URMs in the professoriate may ultimately lead to increased amounts of role models and mentors for the next generation of URM scientists and professors, which will likely have a positive impact on the recruitment and retention of a diverse STEM workforce (5–8).

In several ways, discipline-focused scientific societies are well-positioned to catalyze these interventions and support the professional development of URM researchers. Unlike individual degree programs or postdoctoral positions, scientific societies provide longitudinally stable touchstones for individual scientists across their training and career trajectories, from the undergraduate to senior faculty stages. Societies often assist scientists in obtaining admission to graduate programs or postdoctoral positions, writing grant proposals, publishing research, and presenting their work through poster and oral presentations (9). Unlike individual universities, societies help unite geographically distant scientists and can promote cross-institutional collaborations, networking, mentoring, and career opportunities, benefits that may be particularly impactful for women and URM researchers (9). Given these characteristics, scientific societies can be considered communities of practice (C of P), gathering individuals with similar interests to learn from one another and improve their fields (10). It is often the case that these C of P have explicit or implicit requirements for *bona fide* membership, such as evidenced mastery of a set of skills, abilities, and accomplishments (11). And, although societies can sometimes be inequitable or unwelcoming environments for women and URM researchers (10–15), they are also capable of remedying these problems themselves (16) and can serve as valuable, inclusive, international, professional and social networks for scientists with common intellectual interests (13, 15).

Professional societies can facilitate the integration of URM scientists into their C of P through the creation and implementation of programs to promote the diversity and inclusiveness of their membership and the STEM academic workforce. Indeed, a recent public Request for Information by the National Institute of General Medical Sciences (NIGMS), soliciting “strategies to enhance postdoctoral career transitions to promote faculty diversity, specifically in research-intensive institutions,” revealed several unmet needs and proposed solutions that are well-suited to interventions by societies (17–18). For example, respondents frequently cited bias and inadequate mentoring in academic environments as two major impediments to URM scientist advancement into the professoriate, and proposed solutions such as enhanced mentoring and networking opportunities, inclusive and welcoming environments, and skill development, including grant-writing and faculty job search preparation (17–18). For decades, scientific societies have leveraged their unique strengths to develop and administer a range of programs that promote diversity, inclusion and equity in the biosciences. Many of the publications about these scientific society programs are descriptive in nature, mostly assessing short-term impact on trainees, while few address the outcomes of long-term engagement of trainees with their professional organizations.

In this article, we aim to fill this gap by describing the synergistic ways in which the American Society for Cell Biology (ASCB) and its Minorities Affairs Committee (MAC) have met the challenge of diversifying the biomedical professoriate through programming that addresses some of the most common deficits encountered by URM individuals wanting to join scientific C of P. We also discuss data collected by our organization indicating the benefits and challenges that URM trainees experience during long-term scientific society engagement. We find evidence that increased representation of URM scientists is obtainable through the work of scientific societies. We advocate for scientific societies not only to engage in specific programming to increase the inclusion of URM scientists, but also to assess the outcomes and efficacy of this programming to facilitate evidence-based modifications and improvements. The results of these assessments can shape the future of our scientific societies and how their diversity-related missions are addressed. We also discuss below how the ASCB, in part informed by the evidence presented, is currently working to shift its culture by bringing diversity into the purview of a wider range of members and governing committees, beyond the MAC.

## DISCUSSION

The ASCB, an international professional society of cell biologists founded in 1960, is “dedicated to advancing scientific discovery, advocating sound research policies, improving education, promoting professional development, and increasing diversity in the scientific workforce” (19–20, https://www.ascb.org/about-ascb/). Since 1980, the ASCB has relied upon its MAC “to significantly increase the involvement of underrepresented minority scientists in all aspects of the Society, promoting equity and a sense of belonging for URM cell biologists across their career trajectories” (19–20, https://www.ascb.org/committee/minorities-affairs/). Thanks largely to MAC efforts, ASCB was awarded a 2004 Presidential Award for Excellence in Mathematics, Science, and Engineering Mentoring, the highest honor bestowed upon mentors who work to expand STEM talent (http://paesmem.net/node/1765). Early on in its history, the MAC helped Society leadership integrate diversity and inclusion goals into the ASCB mission, established working relationships with URM-focused committees at other societies and successfully competed for funding from the National Institute for General Medical Sciences (NIGMS) to support specific programming, beginning a tradition that has continued to the present day in the form of support from the NSF and the National Institutes of Health (19–20). The MAC has provided educational and career development opportunities for URM trainees for 30 years (19–20, L. Hammonds-Odie, M.J. Leibowitz, and M. Zavala, presented at the 2017 TWD Program Directors’ Meeting, Baltimore, MD, 18 to 21 June 2017), during which MAC members have used their multidisciplinary scientific and pedagogical expertise to mentor and inspire the next generation of cell biologists from URM backgrounds at a variety of academic stages, from undergraduate/graduate students and postdocs to junior faculty. ASCB MAC initiatives have included programs aiming to increase the success of URM trainees in the science faculty workforce (Table 1). We describe these programs briefly below to set the stage for a discussion of the long-term outcomes associated with involvement in these MAC programs, as reported by trainees in our study.

**Table 1.** ASCB MAC program offerings and percentage/number of respondent participants. ASCB MAC Programs described in this article are listed in this table alongside the percentages and numbers (in parenthesis) of respondents participating in each offering. Program offerings are listed in decreasing number of respondent participants. 54% of respondents indicated having been participants in at least 2-5 ASCB MAC program offerings. The Junior Faculty and Postdoctoral Fellows Career Development Workshop, reporting the highest percentage, allowed for cohort sizes of at least 30 was the activity in which most of the respondents participated.

| **ASCB MAC Program Offering** | **Percentage (#) of Respondent Participants in Program** |
| --- | --- |
| **Junior Faculty/Postdoctoral Fellows Career Development Workshop** | 71% (70) |
| **MAC Travel Awards to ASCB Annual Meeting** | 56% (55) |
| **Visiting Professor (VP) Program** | 31% (30) |
| **Linkage Fellows (LF) Program** | 19% (19) |
| **Marine Biological Laboratory at Woods Hole, MA** | 6% (6) |
| **MAC Mentoring Academy** | 4% (4) |
| **Faculty Research and Education Development (FRED) Program** | 2% (2) |

### Junior Faculty/Postdoctoral Fellows Career Development Workshop

The MAC’s Junior Faculty and Postdoctoral Fellow Career Development Workshop was held annually during the summer months, for about 30 attendees per year, from 2006 through 2017 (20). The 2-3-day workshops aimed to help trainees develop the skills needed for success as they transitioned into tenure-track positions. The workshops included sessions on challenges and opportunities for URM populations, goal-setting and time management, getting published, mentorship, professional conduct, laboratory management and career development at research and teaching institutions, getting the job, NSF and NIH grant opportunities, securing tenure and career advancement, collaborations, networking, and getting started as a new investigator. The workshop curriculum was revised over time to fit the needs of the trainees. Early on, the workshop included opportunities for interested attendees to submit grants or manuscripts in advance for review and feedback from workshop and ASCB faculty. More recently, the MAC deployed new methods to advance diversity and inclusion goals, including adapting an applied theater approach to build skills in conflict resolution and managing difficult professional interactions for female and URM postdocs and junior faculty (21). Ultimately, some of the elements of this workshop were earmarked as ideas for seeding new programming. For example, the ASCB MAC later sought to secure funding and establish a new program focused on participants submitting grants for feedback from experienced mentors—the **Faculty Research and Education Development (FRED) Program,** described in detail below.

### MAC Travel Awards to ASCB Annual Meeting

ASCB MAC Travel Awards enabled approximately 70 URM students and scientists per year, including students and faculty from minority-serving institutions (MSIs), to attend the annual Society meeting in December. Trainees who applied for awards were required to submit abstracts and present posters at the MAC/Education Committee-organized poster competition. For faculty members, preference for travel awards was given to those who were early in their careers and who planned to bring students to the annual ASCB meeting. To support these awardees, the ASCB MAC provided professional development sessions at the annual meeting that awardees were encouraged to attend. These sessions were open to all ASCB members, giving both URM and WR scientists an opportunity to receive mentoring, exchange perspectives, and network with other scientists in similar career stages.

### Visiting Professor (VP) Program

The VP program, which began in 1997, matched over 80 junior faculty members from UR backgrounds and/or MSIs with host senior investigators at research-intensive institutions for one or two summers (8-10 weeks) of research in the senior investigator’s laboratory, with the intent of strengthening the research and educational activities at the VPs’ home institutions. An optional second year of funding was part of this program in later years and allowed the research relationship to continue and for the results of the collaboration to be published. The MAC has determined that participation in the program resulted in more publications and successful grant awards to the VPs, compared to a control group of peers (22).

### Linkage Fellow (LF) Program

The LF program was established in 2000 to provide over 30 faculty members at MSIs with funding for outreach activities supporting cell biology and science career development of students at their home institutions. The LF program has supported workshops, undergraduate research experiences, and outreach to local middle and high schools (20). One of the main objectives of this program was to promote the activities of the MAC and the ASCB at colleges and universities that might not be aware of such activities and to recruit faculty and students to attend the annual meetings.

### Participation in Marine Biological Laboratory (MBL) training

The ASCB MAC provided funding to over 35 individuals from UR backgrounds and/or MSIs to attend scientific courses at the MBL. In this way, the MAC enabled trainees to develop the technical skills needed for their academic success.

### Mentoring Academy

The Mentoring Academy was developed in collaboration with ASCB’s Committee for Postdocs and Students (COMPASS) and targeted postdoctoral fellows at major research institutions. The Mentoring Academy included mandatory training on proactively managing mentoring relationships, which are critical for academic and professional success (23).

### Faculty Research and Education Development (FRED) Program

This program began in 2014 and has bolstered grant development skills for 36 UR and/or MSI postdocs and junior tenure-track faculty members. This was accomplished through the establishment of structured mentoring relationships focused on grant development. FRED fellows were paired with more senior, well-funded and established scientists for a oneyear, grant writing/submission one-on-one mentoring relationship. The program started with a three-day workshop that included grant writing sessions and presentations by both junior and senior faculty and postdoctoral fellows. So far, over half of the participants have secured funding from the NIH, NSF, or other agencies.

### MAC Program Short-term Outcomes: End-of-Program Surveys

The success of the programs outlined above was regularly assessed by anonymous surveys administered to participants by an external evaluator. The majority of participants consistently reported positive objective outcomes and specific benefits of MAC programs, including enhanced networking, successful grant applications, promotions and other professional advancement, publications, research collaborations and curricular practices (21–22).

### MAC Program Long-term Outcomes: Retrospective Survey Analysis

To better understand the long-term outcomes of trainee participation in ASCB MAC programs, we recently surveyed postdoctoral fellows and junior faculty who participated between 2006 through 2015, with 98 (46%) of 215 individuals completing at least part of the survey (see Appendix 1 for study design and methods; approved by Institutional Review Board at High Point University; protocol number 201811-755). Interestingly, most respondents participated in more than one MAC program (54%) and most participated in a program more than three years ago (79%), highlighting the potential of these interventions to fill the needs of trainees and have a positive impact on their career trajectories. The percentages and numbers of respondents participating in each of the MAC program offerings are listed on Table 1 (see also Appendix 2 for demographic and descriptive information of survey respondents). Participants have progressed in their careers, with 88% continuing in academic careers involving some combination of teaching, research, and/or administration (Table 2). 79% credited MAC programs with helping them attain academic promotions, 63% serve as PI or Co-PI on research grants, and 52% have published scholarly articles or book chapters since their participation (Table 3). 53% credited MAC program participation with receiving awards, honors, or other distinctions (Table 3).

**Table 2.** Career Stage at time of first ASCB MAC experience and now. Percentages and numbers (in parenthesis) of respondents disclosing their career stage for “then” (at the time of first MAC program involvement) and “now” (current) categories.

| **Career Stage** | **Then** | **Now** |
| --- | --- | --- |
| Graduate Student | 16% (16) | 0% (0) |
| Postdoctoral Fellow | 31% (31) | 1% (1) |
| Assistant Professor | 43% (43) | 24% (24) |
| Associate Professor | 6% (6) | 41% (41) |
| Full Professor | 0% (0) | 14% (14) |
| University Researcher | 0% (0) | 4% (4) |
| University Administrator | 0% (0) | 4% (4) |
| Private Sector | 0% (0) | 3% (3) |
| Other | 2% (2) | 7% (7) |

**Table 3.** Respondents crediting MAC involvement in their career success.

| Area of Career Success | Respondents |
| --- | --- |
|  | % (n) |
| Promotions | 79% (57 of 76) |
| Presenting at conferences | 76% (56 of 74) |
| Tenure | 67% (34 of 51) |
| Winning grants | 63% (44 of 70) |
| Taking on leadership roles | 63% (49 of 78) |
| Serving on committees | 57% (42 of 74) |
| Receiving awards, honors or other distinctions | 53% (37 of 70) |
| Authoring journal articles and/or other scientific publications | 52% (39 of 75) |

Moreover, past trainees credited MAC programs for the attainment of psychosocial benefits, such as improved selfefficacy (Table 4). 90% of respondents reported that MAC programs helped them feel a part of a scientific C of P and looked forward to continued interactions with networks developed through MAC programs (Table 5). This result highlights the potential of scientific societies as an on-ramp catalyzing membership to scientific Cs of P.

**Table 4.** Respondents credit ASCB MAC programs for the attainment of intangible benefits such as the development of specific soft skills.

| Area/Intangible Benefit | Respondents |
| --- | --- |
|  | % (n) |
| Setting an example for others | 81% (66 of 81) |
| A sense of belonging to a scientific community | 78% (65 of 83) |
| Identity as a scientist | 75% (62 of 83) |
| Feeling fulfilled in a chosen career | 74% (62 of 84) |
| Intention to continue in chosen career | 73% (60 of 82) |
| A sense of self-efficacy | 71% (59 of 83) |

**Table 5.** Contribution of MAC program participation to being part of a “Community of Practice”.

| “C of P” Statement | Respondents |
| --- | --- |
|  | % (n) |
| I feel that I am part of a scientific community of practice, due in part to my involvement in ASCB MAC programs. | 89% (73 of 82) |
| Participation in MAC programs has helped strengthen my identifying as part of a scientific community of practice. | 88% (72 of 82) |
| I look forward to interacting with scientific colleagues met through MAC programs. | 91% (73 of 80) |

### Building on our past successes to amplify impact

The MAC, in part informed by the evidence presented above, is currently working to shift the culture of ASCB by integrating diversity matters into a wider range of activities for members and governing committees, beyond the MAC. We hypothesize that by including WR ASCB members and leadership in these programs, we will augment the power of our interventions for UR, ultimately benefiting all members. Examples include new initiatives like an Prize for Excellence in Inclusivity Award that is one of the featured annual meeting awards (sponsored by the Howard Hughes Medical Institute, https://www.ascb.org/award/new-ascb-prize-for-excellence-in-inclusivity/), a keynote talk and poster sessions at the ASCB annual meeting, focused on the Scholarship of Diversity (https://www.ascb.org/meetings/scholarship-of-diversity/), diversity training for ASCB leadership and Council members, as well as a revamped workshop for postdocs and junior faculty that is not limited to a summer, but which continues for two years and maximizes the use of other annual meeting activities (https://www.ascb.org/career-development/2020-accomplishing-career-transitions-act-program/). Most of these new efforts are currently funded by a NIH-NIGMS IPERT grant (https://www.ascb.org/careers/ascb-receives-nigms-ipert-fundingdiversity-career-development-programs/). By bringing diversity and inclusion to the forefront of ASCB annual meeting programming, we aim to amplify the impact of our efforts and accelerate progress on diversifying the professoriate.

## CONCLUSION

To achieve a more inclusive and thus higher-performing STEM workforce and professoriate, we must recognize that while URMs are well prepared to contribute to the scientific endeavor at the professoriate level (as evidenced by increasing amounts of URMs that are NIH-supported Ph.D. recipients), they may lack the cultural capital needed to succeed in our hyper-competitive academic landscape (3). Scientific societies can provide UR scientists with the training and mentoring needed to meet the challenges of academic professions. In fact, while years of expansive programming have increased representation in a variety of career stages of the STEM workforce, attainment of tenured faculty positions remains a tenuous transition that should be the focus of particular attention (2, 24–26).

We have outlined examples of how such broad challenges can be addressed through programs within scientific societies. Unlike individual institutions, scientific societies are nationwide networks offering discipline-rooted professional support and leadership opportunities. Because the leaders of scientific societies are often standard-bearers in their fields, they have the credibility and experience to serve as excellent role models to younger generations of professionals. Scientific societies owe it to their membership to be as inclusive as possible and to help train future generations of scientists and academicians. Fully diversifying the STEM workforce calls for scientific societies to continue and expand programming designed to increase the URM proportion of undergraduate degree recipients and faculty in particular. The programs and data reviewed here suggest that increases in URM scientist representation at these levels are obtainable through the work of scientific societies.

We advocate for more scientific societies to engage in specific programming to increase the proportions of URM scientists at the bachelor’s and faculty stages. Since many best practices, model programs, and Cs of P have already been established, societies can continue this work and develop additional, targeted programs either alone or in collaboration with one other. It is our hope that, in this way, scientific societies will further increase URM participation at key career transitions, helping to achieve a truly inclusive and productive future STEM workforce.

Moving forward, creating and implementing successful, evidence-based interventions to broaden participation by UR scientists in the biomedical enterprise will likely require not only systemic approaches across the career trajectory, but also centralized collection, analysis and reporting of participant data to guide future programming (1, 4, 12). As trans-institutional organizations that foster the professional development of individual scientists across a wide range of career stages, scientific societies are well-suited to help drive forward the diversification of the academic STEM workforce. ASCB looks forward to working with and learning from other societies in our collective efforts to develop, evaluate and optimize programs that broaden participation in the biomedical workforce.

## SUPPLEMENTAL MATERIALS

Appendix 1: Study Design and Methods Summary

Appendix 2: Demographic Information of Respondents

## ACKNOWLEDGMENTS

The authors thank the ASCB for the Project Initiation Funds that allowed for the collection of the data discussed in this publication. We also thank Joy Quill (https://quillassociates.net/) for serving as an external evaluator of MAC programs. We appreciate the assistance of Irelene Ricks, Deborah McCall, Desiree Salazar, Sara Volk de Garcia, and Fabiola Chacon in implementing and managing the MAC programs mentioned in this publication. We especially thank past participants of ASCB MAC programs.

## Appendix 1: Study Design and Methods Summary

The MAC carried out a long-term outcomes survey to address the following research questions:

1. What impact has MAC program experience had on participants’ career progress, as gauged by publications, grants, promotions, tenure, and similar indicators of career success?
2. To what extent has involvement in MAC programs resulted in intangible benefits and contributed to a sense of belonging to a community of practice (C of P)?
3. Does involvement in more than one program, over a period of years, reinforce the impact on participants’ careers?
4. Does positive impact, if any, persist even after participant involvement comes to an end?

Between 2006 and 2015, approximately 300 graduate students, postdoctoral fellows, and junior faculty members from underrepresented groups participated in career development activities organized by the ASCB MAC. Current email addresses for 215 of these participants were identified. These 215 individuals received a solicitation to respond to the survey that was approved by the High Point University Institutional Review Board (protocol number 201811-755). The survey was administered electronically using the ASCB MAC’s Survey Monkey account. 98 individuals completed at least part of the survey (45.6% response rate) in January 2019.

## Appendix 2: Demographic Information of Respondents

Of the 98 respondents, over half were female (Supplemental Table 1, 58%) and one-third (Supplemental Table 2, 32%) self-identified as Hispanic or Latino. Over 30 of the survey respondents did not answer the question on race (Supplemental Table 3). Of the 68 who did answer, over half (Supplemental Table 3, 56%) self-identified as African American or Black. 6% (Supplemental Table 3, 4) respondents self-identified as of mixed race (two identified as Black and Caucasian, one as Hispanic and Caucasian, and one as Native American and Caucasian (Supplemental Table 3). No Native Hawaiian, Pacific Islander, or South Asian individuals provided race information (Supplemental Table 3). The respondents also indicated that 29% (Supplemental Table 4, 28) were the first in their family to attend college; 26% (Supplemental Table 4, 25) were not born in the United States; 23% (Supplemental Table 4, 23) had at least one parent born outside of the United States; and 21% (Supplemental Table 4, 21) stated that the primary language spoken at home was not English.

---

**Supplemental Table 1:** Gender of Respondents to MAC Programming Outcomes Survey 2019.

| Gender *n*=83 |  |
| --- | --- |
| Female | 58% (48) |
| Male | 42% (35) |
| TOTAL | 100% (83%) |

**Supplemental Table 2:**
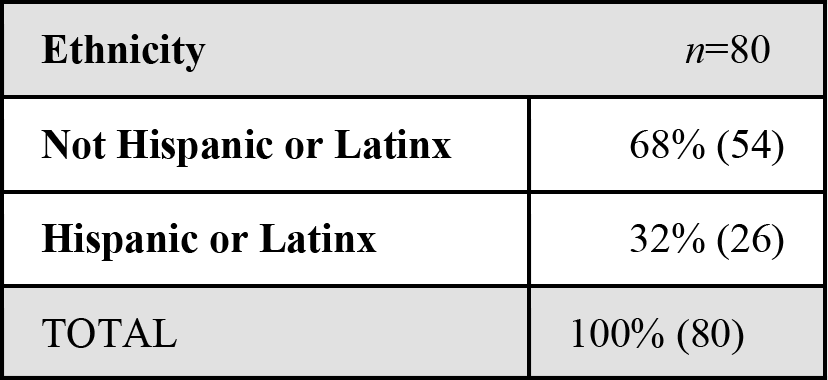
Ethnicity of Respondents to MAC Programming Outcomes Survey 2019.

| Ethnicity *n*=80 |  |
| --- | --- |
| Not Hispanic or Latinx | 68% (54) |
| Hispanic or Latinx | 32% (26) |
| TOTAL | 100% (80) |

**Supplemental Table 3:**
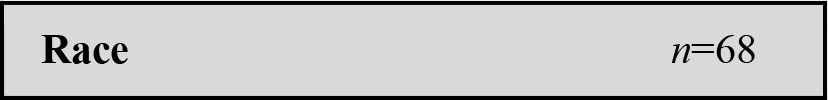
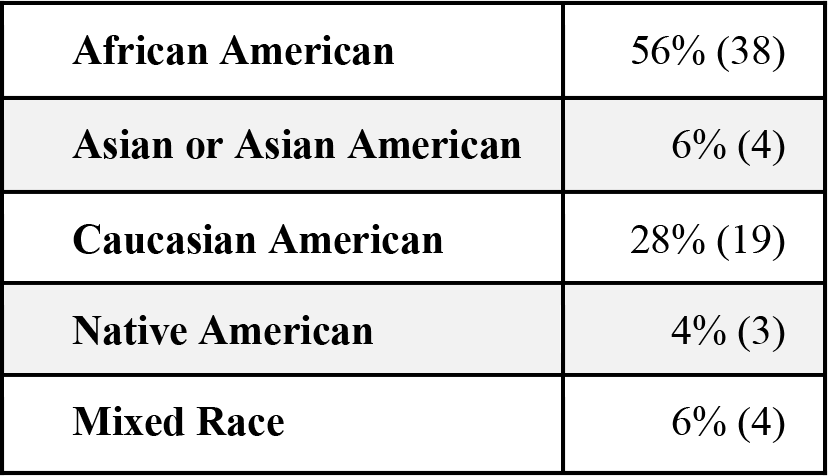
Race of Respondents to MAC Programming Outcomes Survey 2019.

|  |  |
| --- | --- |
| **African American** | 56% (38) |
| **Asian or Asian American** | 6% (4) |
| **Caucasian American** | 28% (19) |
| **Native American** | 4% (3) |
| **Mixed Race** | 6% (4) |

**Supplemental Table 4:** Other Demographic Characteristics of Respondents to MAC Programming Outcomes Survey 2019.

| **Characteristic** | *n*=98 |
| --- | --- |
| **First in family to attend college** | 29% (28) |
| **Not born in the U.S.** | 26% (25) |
| **One or both parents not born in the U.S.** | 23% (23) |
| **Primary language spoken at home is not English** | 21% (21) |

